# An isogenic platform to model the impact of sex chromosome and donor genotype in hiPSC-derived vascular smooth muscle cells

**DOI:** 10.64898/2026.11.18.767725

**Authors:** Heather R. Glatt-Deeley, Isaac V. Faustino, Maria K. Armillei, Erin C. Banda, Darcy T. Ahern, Kevin M. Child, Stefan F. Pinter

## Abstract

Monosomy-X results in Turner Syndrome (TS), which beyond its cardinal height and reproductive phenotypes, spans incompletely penetrant cardiovascular phenotypes, including bicuspid valve, hypoplastic left-sided heart syndrome and aortic coarctation. These congenital heart defects (CHD) conspire with hypertension to dilate the ascending aorta, which can progress to rupture and drive early mortality in TS. Genetic contributors to incompletely penetrant phenotypes in TS remain only partially explored, thus complicating risk stratification.

To begin addressing this challenge in donor-specific in-vitro models, we derived monosomy-X human induced pluripotent stem cells (hiPSCs) alongside isogenic euploid control lines from donors mosaic for the second sex chromosome. In embryonic-origin specific vascular smooth muscle cells (vSMCs) derived from these hiPSCs, monosomy-X reveals donor-specific cellular and transcriptomic effects, with expression changes in one donor panel enriched in genes previously implicated in coronary artery disease and ascending aorta diameter. Polygenic risk scores (PGS) for cardiovascular traits mirror this transcriptomic enrichment, and suggest PGS analysis may help to prioritize hiPSC lines for in-vitro modeling of incompletely-penetrant traits related to TS-associated aortopathy.

## INTRODUCTION

Turner syndrome (TS, 1 in ∼2500 live births) results from monosomy-X, and exhibits two cardinal phenotypes: short stature and premature ovarian failure ^1–3^. Yet, additional phenotypes impacting craniofacial, endocrine, and cardiovascular development, are not fully penetrant. Congenital heart defects (CHDs) in TS include bicuspid valve, hypoplastic left-sided heart syndrome and coarctation ^4^. CHDs necessitate cardiological examination and annual or biannual monitoring starting from infancy to address progressive aortic dilation, which can conspire with early-onset hypertension to progress to aortic aneurysms ^5–7^. These events are often fatal, and account for half of the excess risk of circulatory system mortality, including in adolescence to middle age (ages 15-45) ^6^. Despite incomplete penetrance (∼50%) of CHDs in people with TS, life expectancy of the TS cohort as a whole is shortened by over a decade ^8^.

Due to the relative rarity of TS, few genetic contributors to variably penetrant CHDs in TS have been identified ^9^. Although TS aortopathy has recently been associated with synonymous variants in autosomal *TIMP3,* a paralogue of X-linked *TIMP1,* which has been independently replicated ^10–12^, the mechanism by which such variants may increase aortopathy risk remains unclear. Likewise, the developmental mechanisms of variably penetrant TS phenotypes have remained largely unexplored, in part because the monosomy-X mouse model (39,X) fails to replicate any cardinal TS phenotypes or CHDs ^13–15^. This may be attributed to the differing gene content between mouse and human X, particularly in the pseudoautosomal region (PAR) ^16–18^, and suggests that human cell-based models are needed to shed light on the developmental etiology of TS.

As monosomy-X can also arise somatically as a consequence of aging ^19^, we derived X-monosomic and isogenic euploid control human induced pluripotent stem cells (hiPSCs) from aged donors mosaic for the presence of a second sex chromosome ^20^. Because the anatomical locations of craniofacial, left-sided CHD and other variably-penetrant TS phenotypes implicated neural crest cells (NCC), the multipotent progenitors that populate the developing face and heart, we and others ^6^ hypothesized that monosomy-X impaired some aspect of NCC development. We recently used our isogenic hiPSC panels to demonstrate monosomy-X reduces NCC specification, and implicated hyperactive mTOR (mechanistic-target-of-rapamycin) signaling in this process ^21^. Consistent with a compromised neural crest, TS aortopathy appears to reflect the developmental origin of vascular smooth muscle cells (vSMCs), which are the most abundant cell type of the aorta and constitute the tunica media. In the aortic root, vSMCs derive from the secondary heart field, in the ascending and arch aorta from NCCs, and in the descending aorta from the paraxial mesoderm that also gives rise to somites ^22,23^.

Directed differentiation of hiPSCs to embryonic-origin specific vSMCs ^24,25^ has enabled disease modeling of monogenic aortopathy (Marfan syndrome, *FBN1*) ^26^. Here, we followed this methodology to differentiate our X-monosomic and isogenic euploid control hiPSCs from multiple donors into vSMCs, and characterize cellular and transcriptomic phenotypes of monosomy-X in this isogenic context. Our results enable evaluating the developmental fidelity of this model, and suggest that monosomy-X may exacerbate the independent impact of donor genotype on vSMCs differential expression. Because polygenic risk scores (PGS) mirror this donor-specific transcriptomic enrichment, PGS analysis may inform future selection of hiPSC lines for *in vitro* modeling of incompletely-variant cardiovascular phenotypes in TS.

## RESULTS

We differentiated a set of 11 clonal hiPSC lines reprogrammed from three unrelated donors, 10 lines of which were cytogenomically validated and described previously ^20^. We followed the vSMC differentiation protocol via NESTIN^+^ neural progenitors, as previously described ^25,26^ (with minor modifications, see methods), and stained for immunofluorescence (IF) to assess vSMC markers transgelin (TAGLN) and calponin (CNN1) **[Fig. 1]**. The resulting vSMCs were termed neuroectodermal (NE)-vSMCs originally ^25^, and later neural crest (NC)-vSMCs ^26^, which were described to merely require additional passaging of NESTIN^+^ progenitors in identical conditions. Because no markers have been reported to distinguish the resulting terminal vSMCs by their neural vs. mesodermal origin, we stained neural progenitors prior to single-cell passaging into terminal differentiation conditions [**Fig. 1A**].

**Figure 1:**
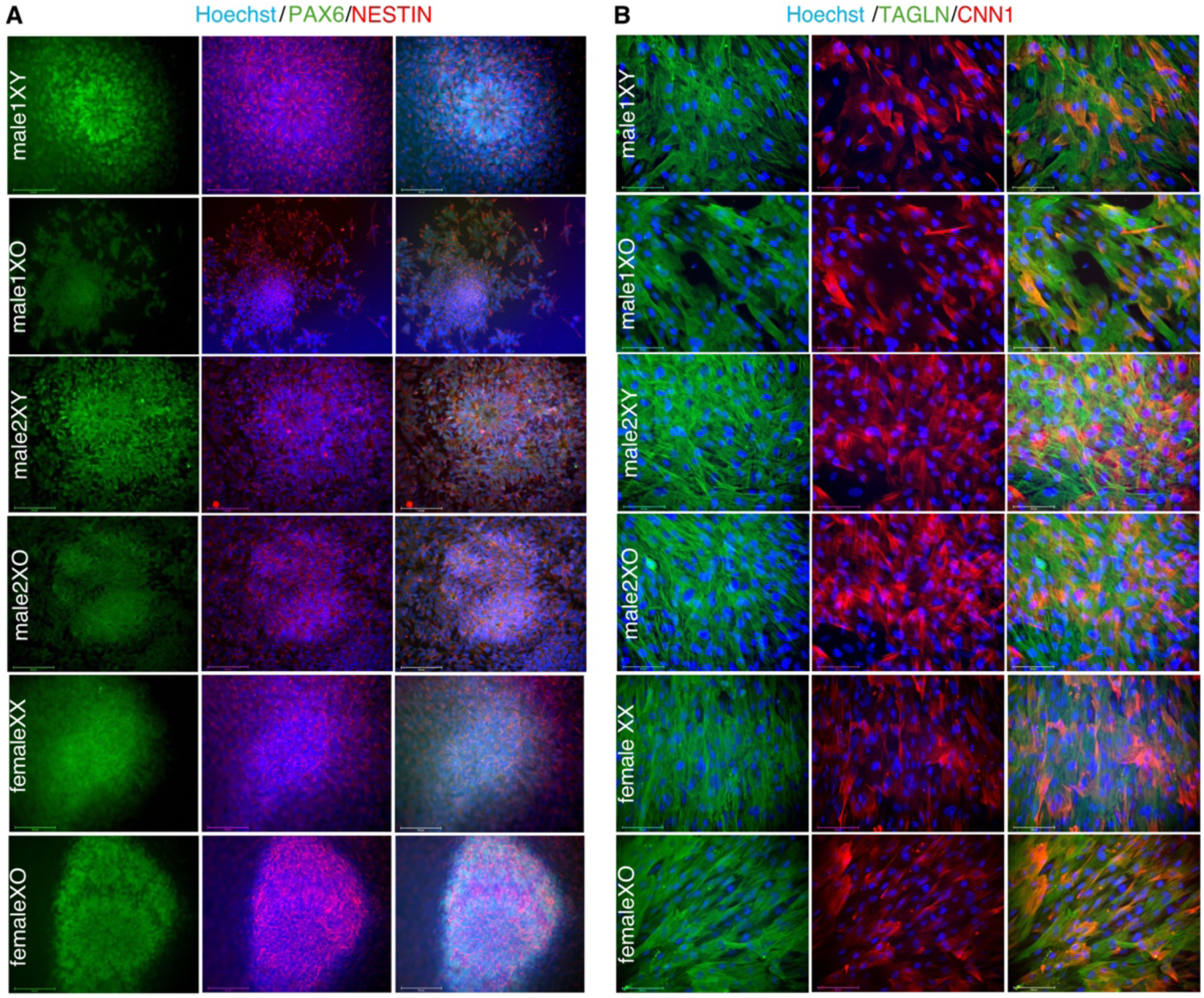
hiPSC-derived vSMCs via NESTIN-positive progenitors. **(A,B)** Representative immunofluorescence (IF) images of hiPSC-derived pairs of monosomy-X alongside euploid control cells stained with **(A)** NESTIN (red) and PAX6 (green) at 6 days of differentiation, or **(B)** CNN1 (red) and TAGLN (green) after 3 weeks of differentiation, following passaging into serum-supplemented media. Channels merged with nuclei (Hoechst33342) and overlaid scale bar (100 µm).

Indeed, we obtained uniformly NESTIN^+^ cells, marking both NC and NE progenitors ^27,28^, while PAX6^+^ cells were observed in neural rosettes, with weak or often no staining in the remainder of the culture. After terminal differentiation in media supplemented by platelet-derived growth factor-BB (PDGFBB) and transforming growth factor-beta 1 (TGFβ1), the resulting cell populations were continually passaged in serum-containing medium, yielding TAGLN^+^ CNN1^+^ double-positive cells **[Fig. 1B]**. This suggests that any heterogeneity of NESTIN^+^ NE (PAX6^+^) and NC (PAX6^-^) progenitor cells may not carry through to the differentiated cell populations.

We next performed mRNA-seq in 3-5 replicates of each clonal line (total of 42 replicates). While validating X-monosomic gene expression, we examined PAR genes and the gametolog gene pairs that were retained on X and Y despite divergent sex chromosome evolution ^29–31^ . As expected, PAR and Y-linked genes were, respectively, reduced or not expressed in X-monosomic samples [**Fig. S1A**]. However, relative to the replicates of the other two male euploid lines, the unpublished male2-derived euploid line featured unexpectedly high expression of all PAR genes, and most Y-linked genes up to and including *NLGN4Y*, as ordered by chromosome coordinates, while *KDM5D* and *EIF1AY* were not expressed [**Fig. S1B**]. This expression pattern is consistent with the presence of isochromosome Y in this otherwise euploid line. We included this line as a separate condition (“m2XYi”) in subsequent experiments, as any differences compared to the other isogenic euploid line may implicate *KDM5D* and *EIF1AY* in monosomy-X associated changes, or conversely rule these genes out. In principal component analysis (PCA) X-monosomic and isogenic euploid controls segregate within each donor, with m2XYi samples clustering close to the other m2XY replicates [**Fig. S1C**]. Likewise, hierarchical cluster segregates samples by donor first, then by karyotype within each donor panel [**Fig. S1D**].

To examine the developmental fidelity of this vSMC model, we compared these expression profiles to recently published human embryonic spatial transcriptome atlases [**Fig. 2**]. First, we assessed markers derived from an early human embryonic cell atlas of early organogenesis (4-6 post-conception weeks (PCW), or Carnegie stages CS12–16) ^32^. Ranking disambiguated gene sets (see methods) of integrated annotations in our vSMC mRNA-seq, the “PA3/4-proximal-2” gene set shows the highest mean expression (y-axis) and is also the most broadly represented [**Fig. 2A**], with >75% of its genes highly expressed (variance-stabilized transformation counts, vst > 10). *In vivo*, migrating NC cells reach the developing heart via pharyngeal arches 3 and 4 to populate the outflow tract ^33^.

**Figure 2:**
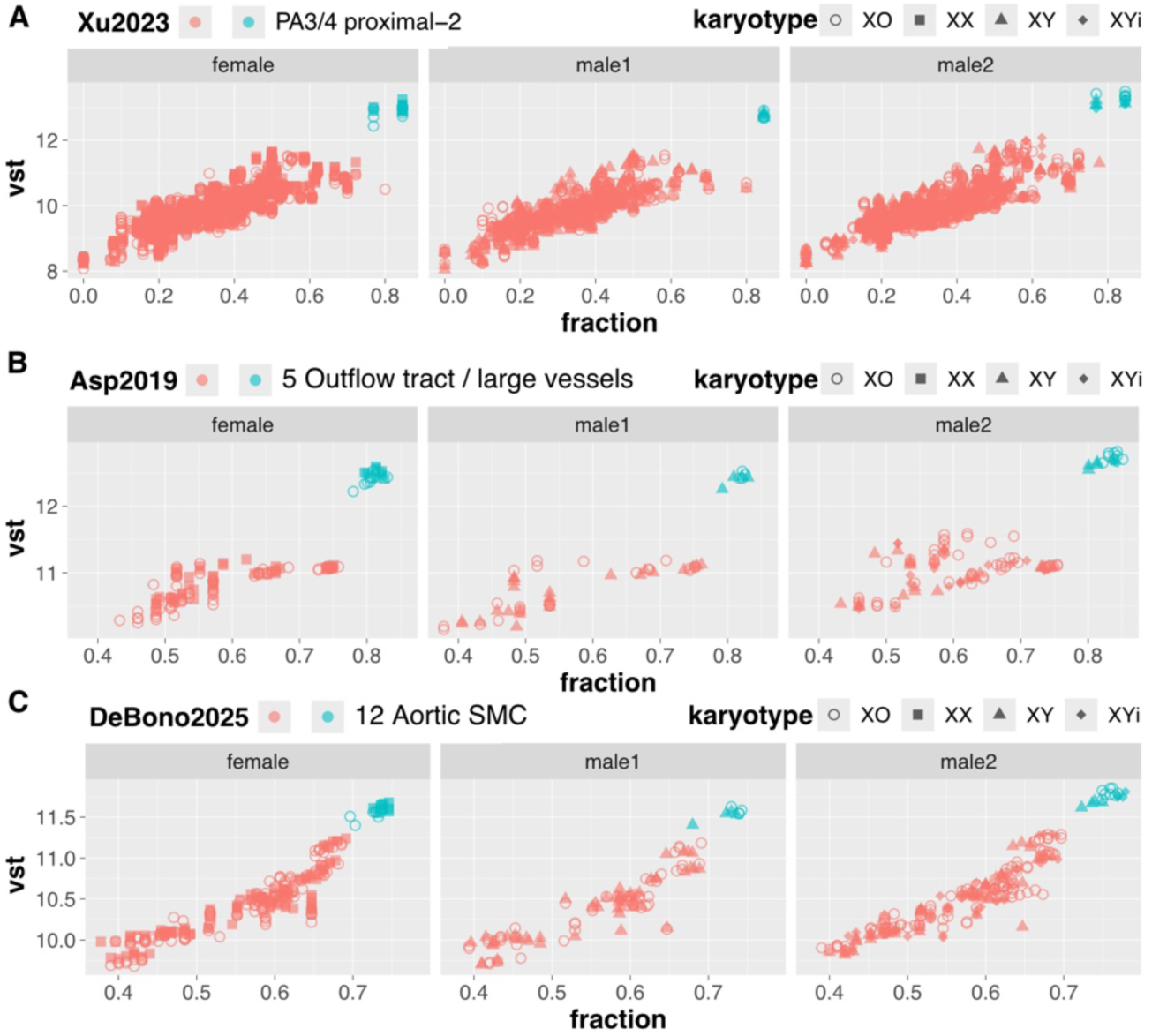
Mean expression of embryonic marker genes annotated by spatial transcriptomes of heart development. Mean expression in bulk vSMC mRNA-seq of annotated genes sets from spatial transcriptomes of embryonic hearts at: **(A)** 4-6 post-conception weeks (PCW) ^32^, **(B)** 6-7 PCW ^34^, and **(C)** 8.6-10.7 PCW ^35^. For each replicate, vst counts are averaged across all genes in each marker set and plotted over the fraction of markers exceeding a mean 10 vst threshold per set. Colored by the most highly expressed marker set in each sample (turquoise) and all others (salmon). Symbols denote karyotype, and faceted by donor.

Consistently, disambiguated gene sets from annotated spatial transcriptomes of CS18 hearts (6.5-7 PCW) ^34^ reveal “Outflow tract / large vessels” genes to be most highly and broadly expressed in our vSMCs [**Fig. 2B**]. Lastly, a higher resolution spatial transcriptome study spanning the prior stages and up to 10.7 PCW ^35^ further ranks the “Aortic SMC” gene set to be the highest and most broadly expressed in our vSMCs [**Fig. 2C**]. In each of the three unrelated donor panels, these marker sets ranked highest in mean expression. Although we cannot assess cell heterogeneity in bulk mRNA-seq, these data suggest our hiPSCs differentiated to TAGLN^+^ CNN^+^ double-positive cells with high and broad expression of genes that would mark NC-derived aortic vSMCs *in vivo*. No overt differences between X-monosomic and isogenic euploid controls in aggregated expression of these gene sets were seen.

Because large-vessel vSMCs have been described to exhibit homeobox (HOX) gene expression dependent on anatomical location ^36,37^, we also compared *HOX* cluster expression. While there were no (female panel) or limited (male1 panel) differences between X-monosomic and isogenic euploid controls, m2XO vSMCs overexpressed several posterior *HOX* genes in all clusters, five of which were shared with m1XO vSMCs **[Fig. S2]**. This is relevant, because forced expression of posterior Hox genes has been shown to impair carotid arteries, aortic arch and descending aorta in the mouse ^38^.

Vascular SMCs have been described to feature wide phenotypic plasticity in health and disease ^39–41^, which also applies to hiPSC-derived vSMCs ^42–44^. We therefore quantified gene signatures associated with contractile and synthetic vSMC states, which were recently reported to correspond to radial position of vSMCs of the great arteries ^45^. Across all three donor panels, we find markers of the synthetic vSMC state to be the most highly expressed, irrespective of whether a subset of “phenotypic modulation” markers is included, followed by the contractile state **[Fig. 3A]**. This is unsurprising, because our vSMC mRNA-seq samples were collected from vSMCs in serum-supplemented media (10% FBS), which is known to promote the proliferative synthetic state in hiPSC-derived vSMCs at the expense of the quiescent contractile state ^46^. We next performed two functional assessments of NC-derived vSMCs, specifically examining their ability to proliferate in TGFβ1-dependent fashion after serum withdrawal, and their contractility. Although we confirmed that vSMCs of all three donors continued proliferating in the presence of TGFβ1 [**Fig. S3**], population doublings clearly slowed relative to the serum-supplemented control, and there were karyotype-dependent changes were not consistent across donor panels. A previous study noted significant donor-specific differences in the response of vSMCs to cytokines ^47^. To assess vSMC contractility, we followed prior methodology ^48^ using CellProfiler image analysis ^49^ of vSMCs visualized before and after treatment with the vasoconstrictor carbachol [**Fig. 3B**]. While contractility of euploid lines somewhat exceeded that of otherwise isogenic X-monosomic cells in female and male1-derived panels, overall contractility lagged somewhat behind what was previously reported for hiPSC-derived vSMCs ^24–26^. In contrast, male2-derived vSMCs contracted more strongly, and X-monosomic cells were hypercontractile relative to isogenic euploid controls [**Fig. 3B**], which was also reproducible with an alternative vasoconstrictor (ET-1, endothelin-1). Importantly, although contractility was assessed in vSMCs cultured in low-serum (0.5% FBS, as in ^46^) media, contractile markers *ACTA2, CALD1, MYH11* and *TALGN* are also elevated in RNA-seq of serum-supplemented (10% FBS) vSMCs from male2, especially in monosomy-X [**Fig. 3A,C**].

**Figure 3:**
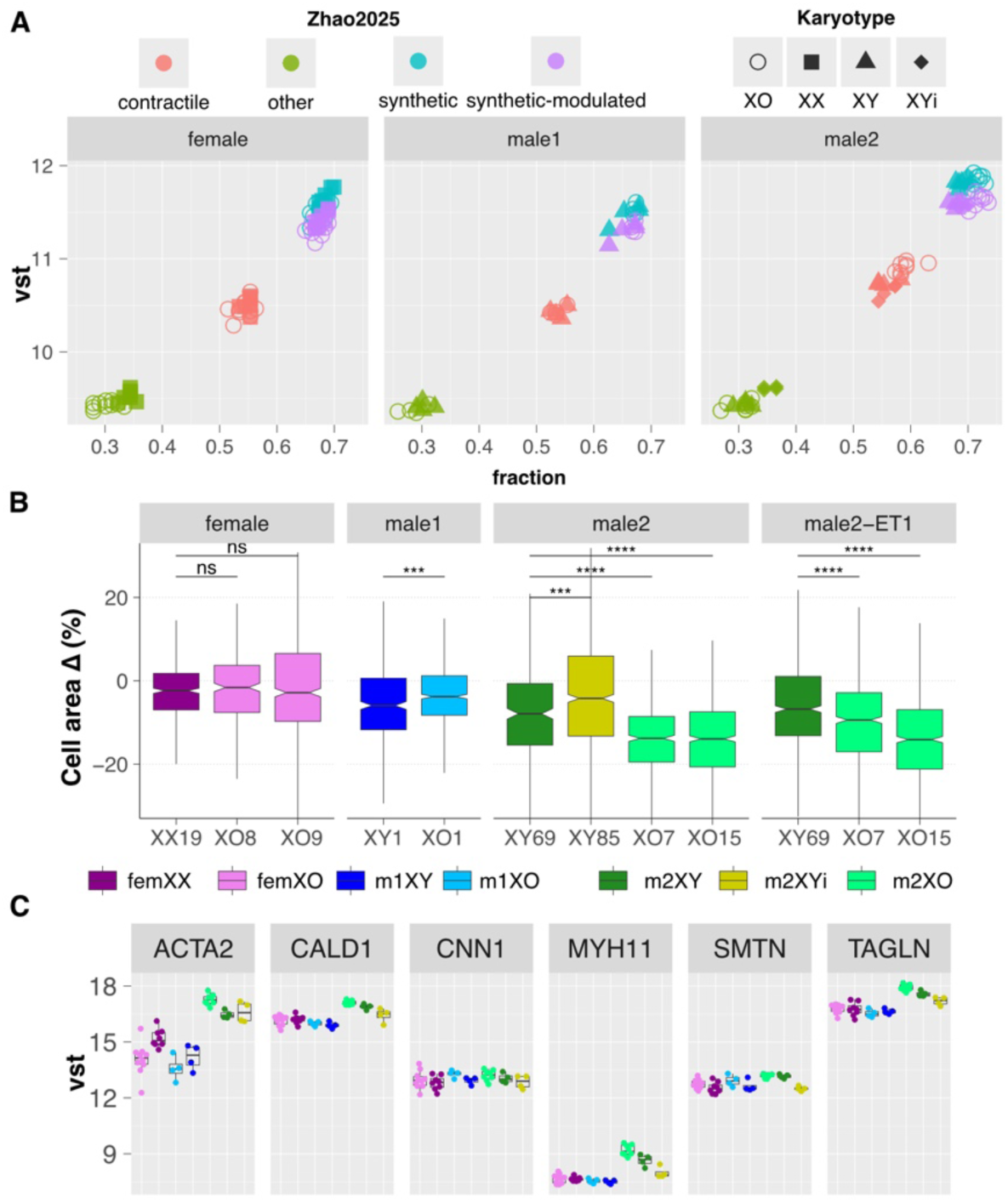
Phenotypic vSMC plasticity reveals male2-specific hypercontractility in monosomy-X. **(A)** Mean expression in bulk vSMC mRNA-seq of annotated genes sets from single-cell transcriptomes of post-mortem human arterial vasculature ^45^, colored by vSMC subcluster with inferred labels: synthetic (subcluster 0, with or without all 19 modulated vSMC genes), contractile vSMC (subcluster 1), and other vSMC (subcluster 2). For each replicate, vst counts are averaged across all genes in each marker set and plotted over the fraction of markers exceeding a mean 10 vst threshold per set. Symbols denote karyotype, and faceted by donor. **(B)** Contractility of vSMCs measured by change in cell area following carbachol treatment (all panels) or endothelin-1 (male2-ET1). Significance (two-sided Mann-Whitney U test) indicated above boxplots (ns, non-significant; ****, p ≤ 0.0001). **(C)** Expression levels (vst) of contractile markers colored by condition.

We next assessed differential expression across the transcriptome. First, we identified differentially expressed genes (DEGs) within each isogenic context, and compared DEGs detected in more than a single donor [**Fig. S4A**]. While a small number of DEGs were common to all three donor panels, 22 of 36 DEGs of which changed in the same direction (concordant, p ≤ 9.5 x 10^-7^), and additional DEGs called in both male1 and female panels DEGs were highly concordant (49/56, p ≤ 5.1×10^−10^), additional DEGs common to male2 and either of the other panels showed only narrowly concordant changes (79/130 with male1, and 86/149 with the female panel). Strikingly, the male2 monosomy-X comparison also detected far more DEGs than the other two isogenic panels combined [**Fig. S4A**], suggesting strong donor2-specific effects may be outweighing concordant changes. Gene-set enrichment analysis (GSEA) of monosomy-X differential expression likewise reveals concordant Reactome gene sets between panels [**Fig. S4B**]. Several of these matched what we previously described in trophoblast and neural crest cell types ^20,21^, including increased cell-cycle and decreased translation gene-sets in monosomy-X, albeit with somewhat reduced concordance in the m2XO comparison.

We then explored alternative transcript isoform usage in isogenic comparisons for monosomy-X within each panel, or donor-specific differences comparing the same karyotype between panels (‘isokaryotypic’). There was little evidence for monosomy-X driven effects in the isogenic comparisons (only three pairwise concordant genes, none triple-concordant). Isokaryotypic comparisons however, indicate that alternative isoform usage is driven by genetics as expected, but again most differential isoform usage involved male2 [**Fig. S5A**]. This male2-specific isoform usage is illustrated by log-transformed P values (logP, cumulative plotted over individual contrasts), colored by the contrast with the highest logP [**Fig. S5B**]. Interestingly, the cumulatively highest-scoring isoform involving male2 [**Fig. S5C**], a differential splice site that lengthens exon 3 of *TOR1AIP1* by a single amino acid, was recently linked to the lead splicing quantitative trait (sQTL) locus associated with coronary artery disease (CAD) ^50^.

Given the preponderance of male2-specific monosomy-X effects, we aimed to understand how these overlapped with monosomy-X differential expression, and whether genes previously linked to cardiovascular disease were enriched in this overlapping set. To this end, we performed GSEA with gene sets identified in individual transcriptome-wide association studies (TWAS) of cardiovascular health. These included genes associated with CAD in the STARNET study ^51,52^ (‘Hao’ and ‘Koplev’), aortic valve stenosis ^53^ (‘Small’), and thoracic aortic aneurysm and dissection ^54^ (‘Klarin’), as well as the diameter of the ascending or descending aorta (‘ascAOR’ and ‘descAOR’), a TS-relevant cardiovascular trait assessed in a landmark UK biobank study ^55^. Most of the significantly enriched gene sets showed higher expression in the male2 panel: these included genes expressed in aorta and associated with CAD (‘Hao’ and ‘Koplev’) ^51,52^ that were enriched in fm2O and mXO2 conditions, respectively **[Fig. S6A]**, as well as genes upregulated in dilated ascending aorta (‘ascAOR+’) ^55^, enriched in mXO1, averaged XO and average male-derived XO (‘ave.XO’ and ‘ave.mXO’) [**Fig. S6B**]. In addition, aortic valve stenosis susceptibility genes (‘Small-’) ^53^ were enriched in all isokaryotypic contrasts involving male2 (‘fm2O’, ‘m1m2O, and ‘m1m2E’) [**Fig. S6C**].

Core genes driving these enrichments are sufficient to cluster conditions and demonstrate the preponderance of male2-specific monosomy-X effects [**Fig. 4**], with 48 DEGs among these 92 genes, all 48 of which are overexpressed in m2XO, compared to only 5 or 6 DEGs in the fXO and m1XO conditions. These genes include high-ranking candidate genes in their respective studies (see ‘ScorePR’, percentiled score band), including *PRDM6* (‘ascAOR+’), *CDKN2B* (‘Hao’), *GALNT2, DCN* and *CXCL14* (‘Koplev’), as well as *PALMD* and *COMP* (‘Small-‘). Although there was surprisingly little gene overlap between studies, with only *PALMD* listed in two enriched core gene sets (‘KoplevAOR’ and ‘Small-‘), we noted cross-study GSEA enrichment of the 15q26.1 CAD-risk locus ^56,57^ containing *MFG8E, HAPLN3* (both ‘HaoAOR’), and *ACAN* (‘Small-‘).

**Figure 4:**
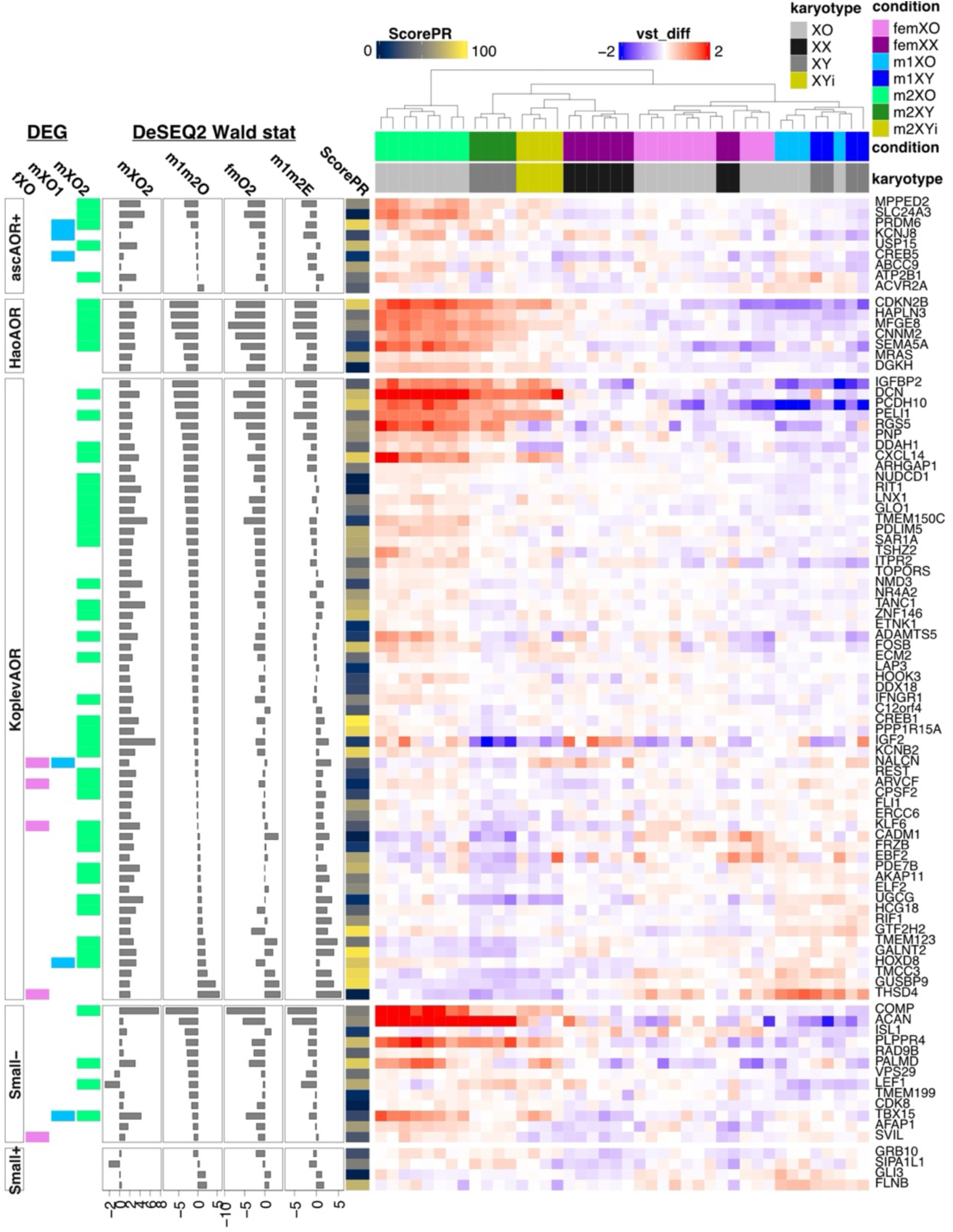
Donor-specific enrichment of genes implicated in cardiovascular disease by transcriptome-wide association. Core genes driving GSEA enrichments of published gene sets associated with cardiovascular traits/disease. Genes grouped by enriched study ^51–53,55^ and study-category (ascAOR+, positively correlated with ascending aorta diameter; HaoAOR and KoplevAOR, CAD driver genes in aorta; Small-/+, expression correlation in aortic stenosis). Samples are clustered by vst differential (sample vst relative to median vst-scaled expression). DEG columns denote differential expression status in monosomy-X relative to isogenic euploid controls. DESeq2 Wald stat columns indicate the differential expression direction and magnitude in isokaryotypic comparisons of female and male1 relative to male2, sorted by the male monosomy-X comparison (‘m1m2O’). ScorePR denotes the percentiled score of each gene within their respective studies.

Because this analysis suggests that monosomy-X exacerbates vSMC differential expression depending on the donors’ genetic background, we wondered whether genotype alone could prioritize cell lines for *in vitro* disease modeling with hiPSC lines from donors without available clinical data. Indeed, prior work has highlighted the predictive value of polygenic (risk) scores (PGS) for informing selection of patient-derived hiPSC lines for *in vitro* modeling of human disease ^58–60^. Because our hiPSC lines were cytogenomically characterized on the CytoSNP-850K array that showed robust imputation and PGS calculation performance across a variety of populations ^61^, we imputed their genotypes against the haplotype reference consortium (HRC) panel, and calculated their PGS (see methods) for a number of cardiovascular diseases and traits, using the Michigan Imputation Server ^62^. Overall, male2 showed significantly lower PGS for cardiovascular disease compared to the female and male1 panels, including for systolic blood pressure, while ranking higher than male1 for diastolic blood pressure [**Fig. 5**]. However, male2 lines ranked first for the ascending aorta diameter PGS (PGS002236 of ∼6 in normalized standard deviations, s.d.), which would translate to an increase in aortic diameter of 3.5 mm in females (effect estimate of 0.59 mm/s.d.), and 4.1 mm in males (0.69 mm/s.d.) in the original validation cohort ^63^. Strikingly, while this GWAS ^63^ and the ascending-aorta TWAS cited above ^55^ both leveraged MRI data from the UK Biobank, our analysis demonstrates methodology-independent concordance between our male2-hiPSC derived vSMC expression data [**Fig. 4**], and male2 PGS [**Fig. 5**]. In sum, our work suggests that PGS analysis may provide predictive power to curate hiPSC panels for *in vitro* modeling of incompletely penetrant cardiovascular phenotypes of TS.

**Figure 5:**
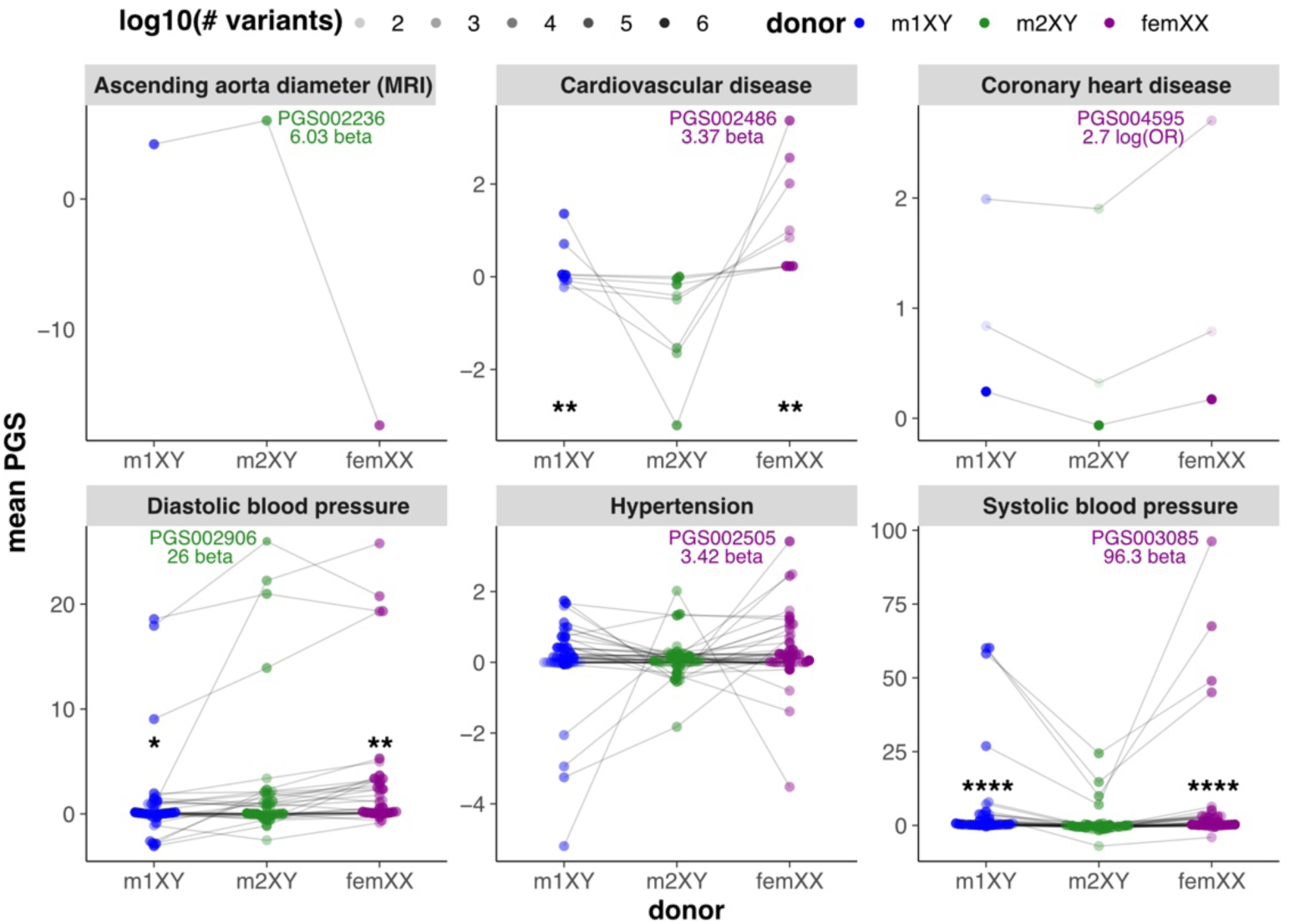
Polygenic risk scores for cardiovascular traits and disease in donor panels. Genotypes imputed from CytoSNP-850K beadchip array calls were used to calculate scores against the PGS catalog on the Michigan Imputation Server. PGS are faceted by Experimental Factor Ontology (EFO) label and grouped by donor colors. Lines connect each individual PGS across donors. PGS opacity reflects the number of variants (log10-transformed) underlying the PGS. Maximal PGS per EFO facet is labeled in the color of the highest-scoring donor panel. Significance of median PGS differences relative to male2 (Mann-Whitney U) indicated above the x-axis.

## DISCUSSION

Several monogenic conditions have revealed that vSMC dysfunction is central to aortopathy ^64,65^, but incomplete penetrance of aortopathy and CHD in TS pose ongoing challenges to risk stratification ^66^. Somatic monosomy-X can also arise with aging due to mosaic loss of the Y (mLOY) in white blood cells, which increases heart ^67^ and renal ^68^ failure, with the latter due to monosomy-X impairing the clearance of senescent cells in the affected tissues. Importantly, mLOY has also been associated with an increased the risk of abdominal, though not thoracic, aortic aneurysms in elderly men ^69,70^, which revealed that mLOY in the aneurysmal aortic tissue itself correlated significantly with both mLOY in T lymphocytes and age ^69^. Understanding how monosomy-X impacts vSMCs is therefore highly relevant to cardiovascular health of women with TS as well as aging men.

To-date, it is unclear why monosomy-X predisposes aortic vSMCs to dysfunction, and whether incompletely-penetrant aortopathy in TS is driven by developmental or genetic variability. Here, we build and validate a platform to begin addressing these questions, using monosomy-X hiPSC lines alongside isogenic euploid controls from three unrelated mosaic donors. We provide an independent assessment of the hiPSC-derived NE/NC vSMC differentiation paradigm ^25,26^, including its reproducibility over 3-5 rounds of differentiation in nearly a dozen hiPSC lines [**Fig. 1**]. Disambiguated marker genes obtained from human embryonic studies are generally consistent with a NC-specific origin of the resulting vSMCs [**Fig. 2**], which most closely resemble aortic vSMCs based on spatial transcriptomics of the developing heart ^34,35^. However, m2XO vSMCs expressed posterior *HOX* genes across all *HOX* clusters [**Fig. S2**], suggesting an altered positional identity that has been shown to impact mouse aorta ^38^. As expected, results confirm the predominant “synthetic” cell state of vSMCs grown in serum [**Fig. 3A, Fig. S3**] with moderately concordant differential expression among panels [**Fig. S4**], but also reveal monosomy-X to impart a donor-specific impact on contractility [**Fig. 3B,C**]. Given similar clustering and contractility of m2XYi with isogenic euploid m2XY vSMCs, it is unlikely that these changes observed in m2XO are due to loss of *KDM5D* and *EIF1AY* [**Fig. 3, Fig. S1**]. Assessing alternative transcript isoform usage [**Fig. S5**], we identified recurrent male2-specific differential splicing of exon 3 of *TOR1AIP1*, suggesting the presence of a lead CAD-associated sQTL ^50^.

Expression of genes associated with heart disease and aortopathy by TWAS was generally elevated in monosomy-X of male2 (m2XO), including some of the top-scoring genes in their respective TWAS studies [**Fig. 4, Fig. S6**]. While such aorta-expressed genes (‘HaoAOR’ and ‘KoplevAOR’) from the STARNET studies of CAD ^51,52^ were significantly enriched by GSEA, there was no clear bias toward overexpression of plausibly protective versus risk-associated genes. Aortic stenosis susceptibility genes ^53^ were also overexpressed in male2 isokaryotypic comparisons, only some of which were further elevated in m2XO vSMCs, including *COMP,* shown to exhibit dynamic expression in aortic valve stenosis ^71^ and aortic dissection ^72^, and previously described *PALMD,* a high-scoring protective factor ^73^. In contrast, only genes whose expression was positively correlated with ascending aorta diameter (‘ascAOR+’) ^55^ were consistently overexpressed in m2XO [**Fig. 4**], including *MPPED2, SLC24A3* and top hit *PRDM6*.

Given the number of male2-specific observations [**Fig. 3,4** and **S4-6**], we were interested to learn whether prior PGS analysis would have prioritized the male2-derived hiPSC panel to model such incompletely-penetrant phenotypes of NC-derived vSMCs. We therefore used our published CytoSNP array data of euploid donor hiPSC lines to impute variants and calculate PGS for cardiovascular traits. Interestingly, relative to female and male1, male2 featured a relatively protective genetic signature with significantly lower PGS for ‘Cardiovascular disease’ and ‘Systolic blood pressure’ [**Fig. 5**].

However, male2 scores higher than male1 for ‘Diastolic blood pressure’, and highest for ‘Ascending aorta diameter’. The latter would imply a 4.1 mm wider aortic diameter over the male average (33.3 mm) in of the validation cohort, reported to feature an elevated hazard ratio of >1.4 for thoracic aneurysm in the UK Biobank ^63^.

Overall, our study suggests monosomy-X impacts hiPSC-derived NE/NC-vSMCs in largely donor-specific fashion. This is consistent with prior reports of primary aortic vSMCs ^47^, and suggests that monosomy-X acts on donor-specific expression to exacerbate phenotypic variability and uncover genetic susceptibility. Importantly, our study demonstrates concordant enrichment of differential expression [**Fig. 4**] and PGS [**Fig. 5**] that would be independently consistent with an increased aortic diameter in male2. As TS aortopathy is incompletely-penetrant, and some genetic predictors like autosomal *TIMP3* have been identified ^10–12^, it is therefore plausible that PGS analysis may be useful in prioritizing X-monosomic hiPSC lines for *in vitro* modeling of TS aortopathy, as it has been for hiPSC modeling of other diseases ^58–60^. Whether PGS may provide a similar predictive utility for risk stratification for TS aortopathy remains to be tested.

While monosomy-X has also been reported to impart subtle congenital ^74^ or induced ^75^ aortic alterations in the X-monosomic mice, the 39,X animal model does not recapitulate TS aortopathy or even the fully-penetrant cardinal TS phenotypes. In addition, the donor-specific impacts reported herein suggest that human cell-based models paired with genetic analysis may prove to be useful towards understanding TS aortopathy. Given genetic, clonal and technical variability, the hiPSC-derived vSMCs model requires multiple donors, clonal lines and replicates that can be challenging to implement and assess in parallel. Prospective PGS analysis in available hiPSC consortia lines, combined with targeted Y chromosome ablation in selected lines ^76^, may therefore provide an especially powerful approach toward understanding TS aortopathy in future studies.

## ACKNOWLEDGEMENTS

We thank Sanjay Sinha’s lab (Cambridge University) and Yibing Qyang’s lab (Yale University) for helpful advice on implementing the vSMC differentiation protocol. We also thank Lixia Yue (UConn Health) for advice on vSMC imaging, and Bo Reese (University of Connecticut, Center for Genome Innovation) for mRNA-seq library preparation and sequencing. This work was supported by NIH grants R01HL141324, R35GM124926 and R35GM153326 to S.F.P.

## MATERIALS AND METHODS

### NE-SMC differentiation

The protocol for differentiating NE-SMC from hiPSCs was as described in ^25^ with minor modifications. Briefly, hiPSCs were grown in 6-well tissue culture plates under feeder-free conditions in mTeSR+ or mTeSR-1 (Stem Cell Technologies), and split (1:4) at ∼80% confluence with 0.5 M EDTA, yielding intermediate clumps of ∼50-100 cells. These were plated on gelatin-coated wells in 2 mL of 1:1 mTeSR+:CDM-PVA medium with 1 µl/mL 10 mM Y-27632 ROCK inhibitor (Fisher Scientific). CDM-PVA medium was made with 1:1 IMDM:F12, and 0.1% poly(vinyl alcohol) (MilliporeSigma) pre-dissolved for 30 minutes at 95°C in a small volume of base media, and supplemented with 15 µg/mL transferrin (MilliporeSigma), 1% chemically defined lipid concentrate (ThermoFisher Scientific), 0.004% monothioglycerol, 1x Pen-Strep, and 7 µg/mL insulin (MilliporeSigma) added after 0.2-micron filtration. To gelatin-coat tissue culture plates, wells were incubated in 0.1% gelatin for at least 15 minutes at 37°C, aspirated, and covered with 1 mL mouse embryonic fibroblast (mEF) media, then used within 1-7 days. Prior to plating cells, media was replaced with CDM-PVA with fresh 12 ng/mL bFGF (ThermoFisher Scientific) and 10 µM SB41542 (MilliporeSigma). On days 3 and 5 (d3, d5), half the media volume in each well was replaced with fresh media (to 2 mL). Overly dense centers of colonies were removed by aspiration, and media changed as necessitated by cell density. On d7, neuroectodermal (NE) cells were gently dissociated and singularized with TrypLE Express (ThermoFisher Scientific) and resuspended in 10x volume CDM-PVA. Cells were pelleted at 210xg, counted, and seeded in 6-well gelatinized plates at approximately 0.5 - 1 x 10^6 cells per well in CDM-PVA with freshly added 10 ng/mL PDGF-BB (Peprotech) and 2 ng/mL TGFβ1 (Peprotech). Except for the first full media change, half-media changes were performed 3-4 times per week until d19 (for CDM-PVA+PT). Confluent wells were split (1:2-3) with TrypLE Express (ThermoFisher Scientific) as needed on gelatinized plates. On d19, the medium was replaced with serum-supplemented DMEM (DMEM, 10% FBS, 1x Pen-Strep, 0.5x Glutamax, all ThermoFisher Scientific except FBS from MilliporeSigma). Full media changes were performed 3-4 times/week. Upon reaching confluency, typically within a week in serum-DMEM, cells were split and replated as before ∼1:2-1:4 on gelatin plates. Cells were considered adapted after approximately a week or more in serum-DMEM and upon having reached confluence.

### Immunofluorescence

Cells were grown in Nunc Lab-Tek II Permanox chamberslides (ThermoFisher Scientific). Cells were fixed in 4% formaldehyde for 10 minutes at room temperature, washed with 0.1% Triton-X and then permeabilized in 1% Triton-X for another 10 minutes. Samples were blocked in blocking buffer containing 0.1% Triton-X, 2% BSA and 5% normal goat serum (MilliporeSigma) at room temperature for one hour. Primary antibody hybridization was performed with NESTIN (611658, 1:500, BD Biosciences) and PAX6 (901302, 1:200, Biolegend) for NE-cells or TAGLN (ab14106, 1:500, abcam) and CNN-1 (C2687, 1:20,000, MilliporeSigma) for SMCs, in blocking buffer overnight at 4°C. Secondary antibodies AlexaFluor555-Goat-anti-Mouse and AlexaFluor647-Goat-anti-Rabbit (1:500, ThermoFisher Scientific) were incubated in blocking buffer for 45 minutes at room temperature. All immunofluorescence slides were washed with 0.1% Triton-X. Nuclei were post-stained before the last wash 2 µg/ml Hoechst 33342. ProLong Gold Antifade Mountant without (for Hoechst 33342-stained slides) or with DAPI (ThermoFisher Scientific) was used to mount slides. Slides were imaged with an EVOS FL Auto system (ThermoFisher Scientific).

### SMC Proliferation assays

NE-SMCs were seeded at 6,000 cells/well into two separate flat-bottomed 96-well gelatin-coated plates in DMEM/F12 without phenol red (ThermoFisher Scientific), and with 1x pen-strep, then incubated overnight. Negative control wells with media but without any cells were included. The next day (d0), the first plate was processed using the CellTiter 96 AQueous Non-Radioactive Cell Proliferation Assay (MTS) (Promega) per manufacturer’s instructions using a Spark Multimode Microplate Reader (Tecan). On the second plate, media was removed and replaced with either DMEM/F12 alone, or with freshly added 10 ng/mL TGFβ1or 10% FBS. Each condition was carried out in at least 6 replicate wells. Media was changed daily on the second plate until d3, when the MTS assay was performed with the second plate as with the first.

### SMC Contractility assays

Serum-adapted NE-SMC were passaged with Accutase (ThermoFisher Scientific) and plated on GelTrex-coated 24-well tissue culture plates in triplicate in DMEM, 0.5% FBS, 1x pen-strep at ∼20,000 cells/well. Cells were fed 3 times per week and assessed for contractility on d7, as described in ^48^. Briefly, cells were stained with 4 µM calcein-AM, red-orange (ThermoFisher Scientific) for 30 minutes, and rinsed twice in DMEM. Following the last rinse, cells were incubated in 0.5 ml DMEM/well for 30 min at 37°C. Once the plate was positioned and calibrated in EVOS FL Autoimaging System, care was taken not to move or bump the plate to avoid changing the position/focus in between images. For each well, a field of view was chosen, manually focused, and a location capture pin created, until up to 6 wells were pinned (near the center of each well). A set of 3 images, each separated by 20 min, was captured for each field of view with the automated scan feature with autofocus selected starting from the first pin. After the first images, the plate cover was carefully removed and 0.5 ml of room temperature DMEM with 20 µM carbachol (MilliporeSigma) or 0.2 µM endothelin-1 (MilliporeSigma) was added to each well for a total volume of 1 mL and final concentrations of 10 µM carbachol (or 0.1 µM endothelin-1). Then the plate cover was carefully placed back on top of the plate. Cells incubated for another 20 min (or 30 minutes for endothelin-1) and the final set of images taken for each pinned field of view.

### CellProfiler analysis

CellProfiler^49^ (version 4.2.4) was used for cell identification, tracking, and area size measurements, based on a previously reported pipeline to SMC contractility measurements ^48^. Briefly, a total of 12 images (6 fields-of-view imaged before and after carbachol/endothelin treatment) was imported into the pipeline at a time, using metadata extraction from the filename to retain clonal line and pre- or post-treatment information. The IdentifyPrimaryObjects, MeasureObjectSize, and TrackObjects modules were respectively used to identify cells within each image, measure their sizes, and add tracking labels to match each cell before and after treatment. Tables reporting object number, metadata, area size, and tracking label across each set of 12 imported images were exported. R (version 4.2.2) was used for statistical analysis and plotting of SMC contractility. Experiments indicating cell object tracking errors across stacks of images due to plates shifting between timepoints were removed from analysis. Individual cells with ambiguous corresponding labels (multiple or missing) in pre- and post-treatment images were removed, as were cells measuring in the top and bottom quintiles for starting cell area size. The final dataset included 6653 identified cells tracked across paired (pre- & post-treatment) images for analysis from 18 different experiments. Using the matched cell area measurements, the absolute and relative (percent change) area difference between pre- and post-drug, and percent of initial area post-drug, were calculated. Median percent change values were calculated for each clonal line used, and compared by two-sided Mann-Whitney U test.

### RNA-seq and PGS analysis

RNA was isolated from serum-adapted NE-SMCs from near-confluent 6-well plates (∼0.7-1 x 10^6 cells/well), with the PureLink RNA Mini kit (ThermoFisher Scientific). Libraries were prepared at the UConn Center for Genome Innovation using the Illumina TruSeq Stranded mRNA Library Prep Kit and 100-bp paired-end reads were sequenced to an average depth of 37.7 (±11.5) million read pairs/replicate on the NovaSeq S6000 (Illumina).

Read pairs were processed for adapter removal with fastp ^77^ and aligned to the human reference transcriptome (GRCh38, GENCODE v.38) using salmon ^78^, followed by differential expression analysis of 17,920/60,230 genes with DESeq2 ^79^, as recently described ^21^. Eight surrogate variables were estimated by sva ^80^, and added to the DESeq2 design that also corrects for round of sequencing and tests for condition, separating karyotypes in donor-specific fashion.

Gene sets from spatial transcriptomic studies across human heart development ^32,34,35^ and post-mortem vasculature ^45^ were assigned to disambiguated celltype/tissue-specific subsets within each corresponding study based on the minimal adjusted p-value (maximally 0,1), while allowing for ties. Genes also required a positive log2FoldChange relative to the other celltypes/tissues within each study for assignment, and each final disambiguated set required a minimum of 10 genes total for inclusion in the analysis. Reactome (MSigDB) gene set enrichment analysis (GSEA) using clusterProfiler ^81^ was performed on all genes ranked by the DESeq2 Wald statistic, and the equal-weighted average of the quantile-normalized Wald scores from euploid-45,X comparisons (fXO, mXO1, and mXO2).

Alternative splicing and transcript isoform usage were assessed using the R package Isoformswitchanalyzer ^82^ using default parameters other than a minimal geneExpressionCutoff of 2. Differential isoform switches identified by IsoformSwitchTestSatuRn required a dIFcutoff of 0.2 (20%) and default FDR-adjusted p-value of ≤ 0.05. Modules analyzeORF, analyzeAlternativeSplicing, and analyzeIntronRetention were run with default parameters.

For PGS analysis, we converted our previously published CytoSNP array data ^20^ from Illumina’s idat to genotype-call (gtc) format, then to variant-call format (vcf) using bcftools ^83^ and the gtc2vcf module of MoCha tool ^84^. Imputation of autosomal variants from these CytoSNP-represented variants was performed on the Michigan imputation server ^62^ against the Haplotype Reference Consortium (HRC) panel ^85^, followed by calculation of polygenic risk scores for cardiovascular disease and measurements provided by the PGS catalogue ^86^.

## Notes

### Competing Interest Statement

The authors have declared no competing interest.

