## Supplement for "An isogenic platform to model the impact of sex chromosome and donor genotype in hiPSC-derived vascular smooth muscle cells"

**Supplementary figures and legends:**

Figure S1: Clonal lines clustered by sex chromosome karyotype and donor.

Figure S2: Ordered HOX cluster gene expression in hiPSC-derived vSMCs.

Figure S3: Proliferation in DMEM and serum or TGFβ1- supplemented media.

Figure S4: Global differential expression and Reactome gene-set enrichment analysis.

Figure S5: Transcript isoform usage in isogenic and isokaryotypic contrasts.

Figure S6: GSEA plots of gene sets associated with cardiovascular disease or traits.

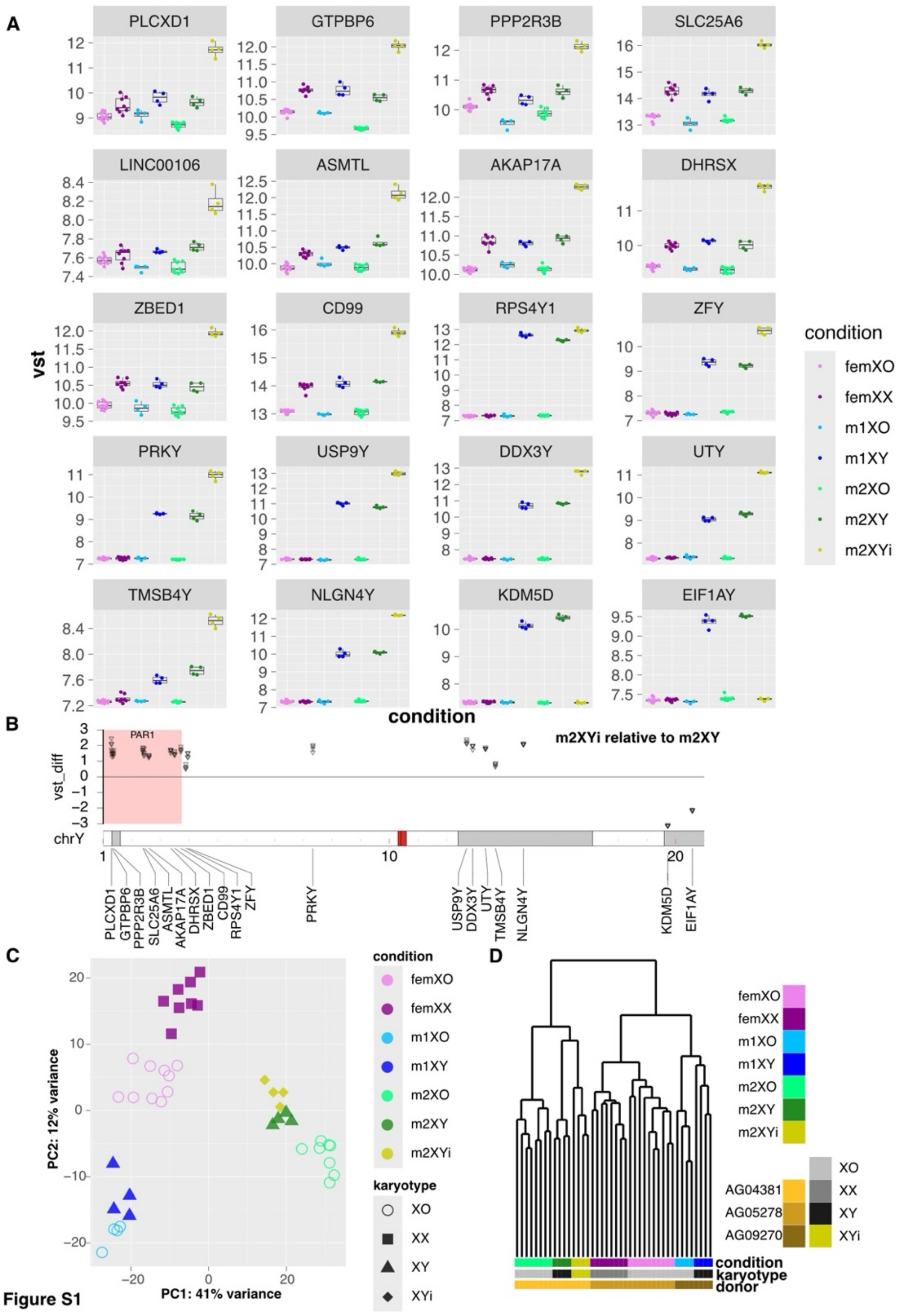
A
PLCXD1
GTPBP6
PPP2R3B
SLC25A6
LINC00106
ASMTL
AKAP17A
DHRSX
ZBED1
CD99
RPS4Y1
ZFY
PRKY
USP9Y
DDX3Y
UTY
TMSB4Y
NLGN4Y
KDM5D
EIF1AY
vst
condition
femXO
femXX
m1XO
m1XY
m2XO
m2XY
m2XYi
B
PAR1
m2XYi relative to m2XY
vst_diff
chrY
C
PC2: 12% variance
PC1: 41% variance
condition
karyotype
XO
XX
XY
XYi
D
AG04381
AG05278
AG09270
condition
karyotype
donor

Figure S1

**Figure S1: Clonal lines clustered by sex chromosome karyotype and donor.** **(A)** Expression levels (vst) of PAR and Y-linked gametolog genes ordered by chromosomal coordinates, and colored by condition. **(B)** Difference in m2XYi expression levels (“vst-diff”) for each gene relative to isogenic euploid controls (m2XY) over the Y (0-21 Mb, PAR shaded in red). **(C)** Principal-component analysis (PCA) segregates samples by ‘‘condition’’ (donor and karyotype), with percent of variance of PC1/PC2. **(D)** Dendrogram of RNA-seq samples segregating by donor and karyotype.

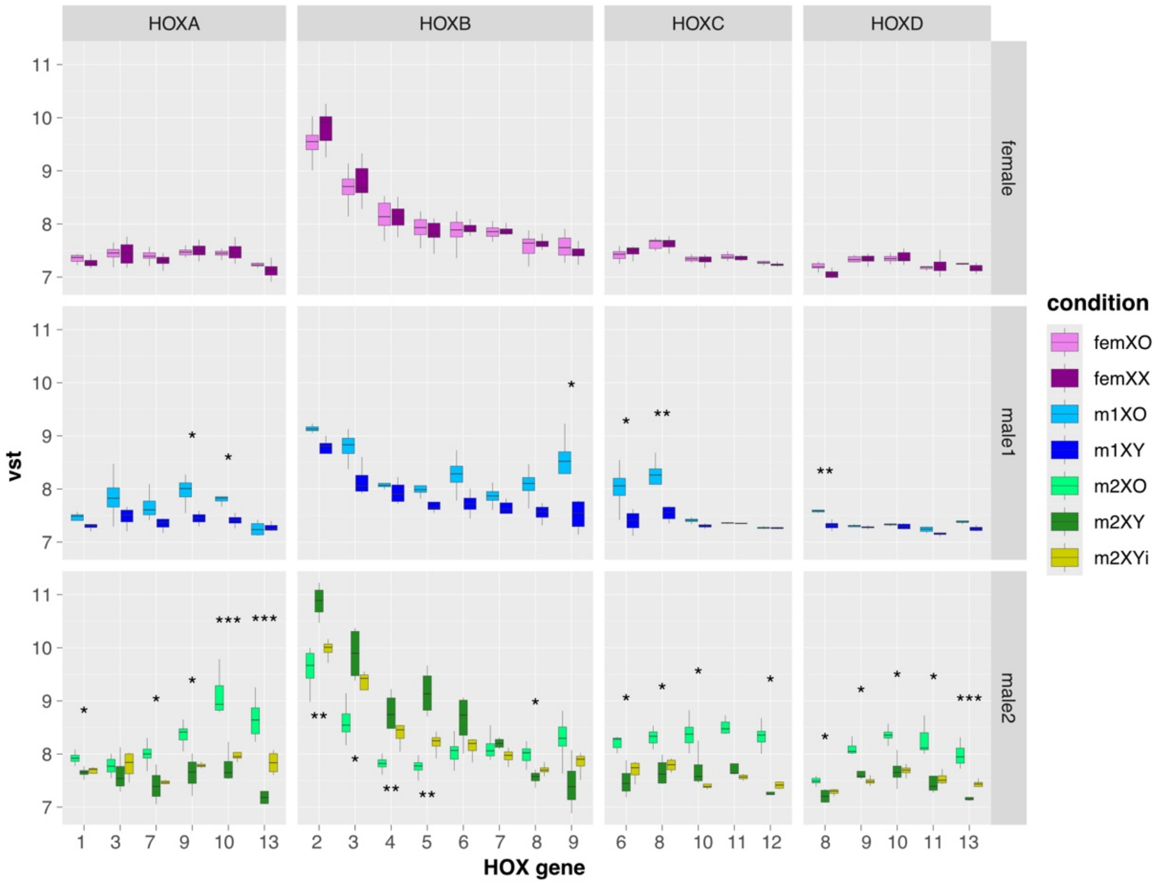


**Figure S2: Ordered HOX cluster gene expression in hiPSC-derived vSMCs.**

Expression levels (vst) of HOX genes as ordered within each HOX cluster, and colored by condition. Adjusted p-value (from DESeq2 Wald test) relative to isogenic euploid controls (*, **, and *** for p.adjust ≤ 0.1, 0.01, and 0.001, respectively) is indicated above (up-regulated) or below (down-regulated) boxplots.

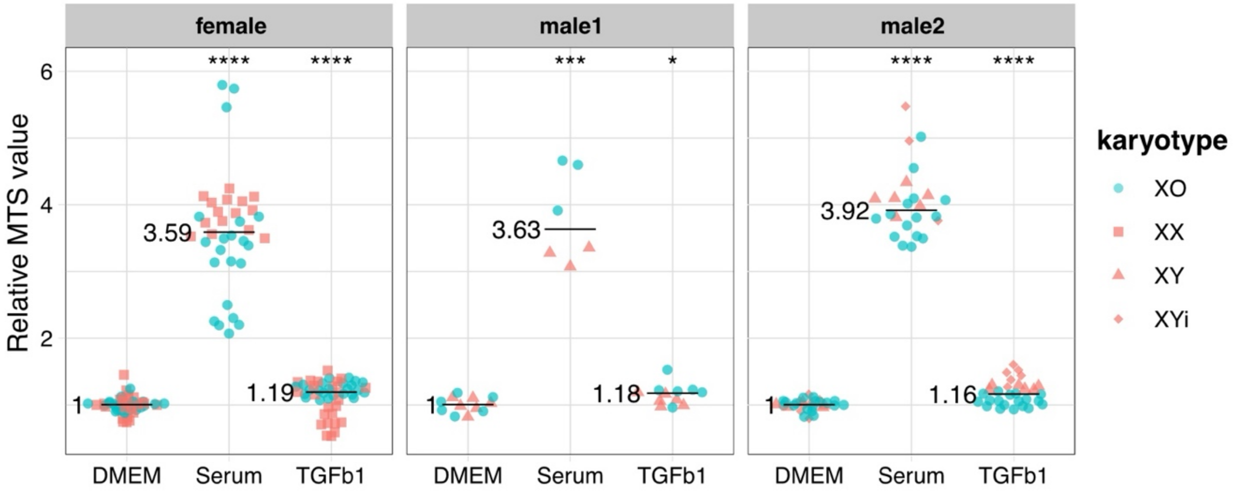


**Figure S3: Proliferation in DMEM and serum or TGFβ1- supplemented media.**

Cell proliferation based on tetrazolium salt (MTS) labeling relative to mean MTS of corresponding DMEM control, as measured after 3 days of culture. Symbols and colors denote karyotype, faceted by donor. Relative MTS median indicated by black crossbar and number label. Significance compared to DMEM control indicated above each plot (two-sided Mann-Whitney U test: * $p \leq 0.05$, ** $p \leq 0.01$, *** $p \leq 0.001$, **** $p \leq 0.0001$).

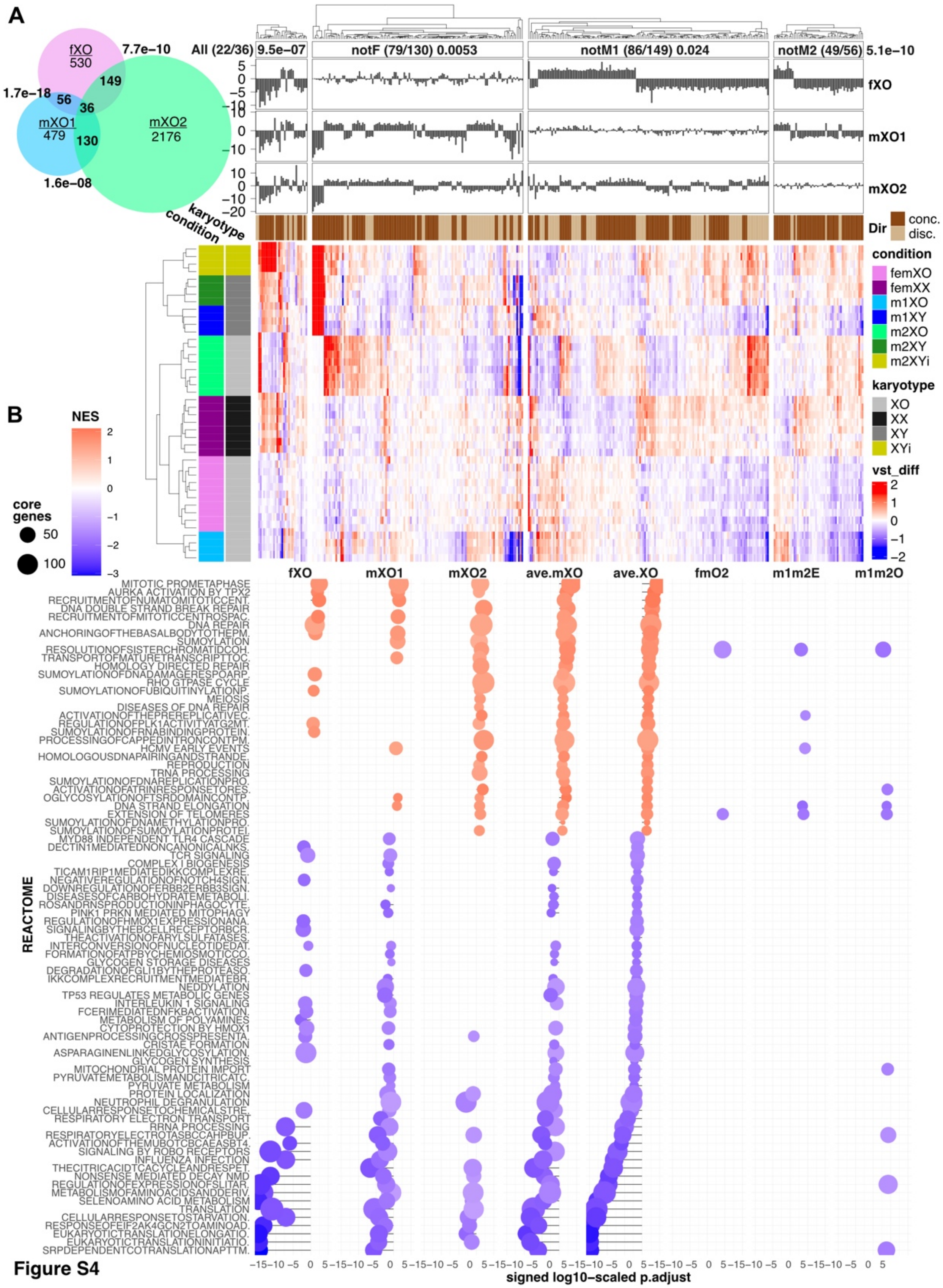
A
fXO
530
7.7e−10
149
1.7e−18
56
36
mXO1
479
130
mXO2
2176
1.6e−08
All (22/36) 9.5e−07
notF (79/130) 0.0053
notM1 (86/149) 0.024
notM2 (49/56) 5.1e−10
fXO
mXO1
mXO2
Dir
conc.
disc.
karyotype
condition
condition
femXO
femXX
m1XO
m1XY
m2XO
m2XY
m2XYi
karyotype
XO
XX
XY
XYi
vst_diff
B
NES
core genes
50
100
fXO
mXO1
mXO2
ave.mXO
ave.XO
fmO2
m1m2E
m1m2O
REACTOME
MITOTIC PROMETAPHASE
AURKA ACTIVATION BY TPX2
RECRUITMENTOFNUMATOMITOTICCENT.
DNA DOUBLE STRAND BREAK REPAIR
RECRUITMENTOFMITOTICCENTROSPAC.
DNA REPAIR
ANCHORINGOFTHEBASALBODYTOTHEPM.
SUMOYLATION
RESOLUTIONOFSISTERCHROMATIDCOH.
TRANSPORTOFMATURETRANSCRIPTTOC.
HOMOLOGY DIRECTED REPAIR
SUMOYLATIONOFDNADAMAGERESPOARP.
RHO GTPASE CYCLE
SUMOYLATIONOFUBIQUITINYLATIONP.
MEIOSIS
DISEASES OF DNA REPAIR
ACTIVATIONOFTHEPREREPLICATIVEC.
REGULATIONOFPLK1ACTIVITYATG2MT.
SUMOYLATIONOFRNABINDINGPROTEIN.
PROCESSINGOFCAPPEDINTRONCONTPM.
HCMV EARLY EVENTS
HOMOLOGOUSDNAPAIRINGANDSTRANDE.
REPRODUCTION
TRNA PROCESSING
SUMOYLATIONOFDNAREPLICATIONPRO.
ACTIVATIONOFATRINRESPONSETORES.
OGLYCOSYLATIONOFTSRDOMAINCONTP.
DNA STRAND ELONGATION
EXTENSION OF TELOMERES
SUMOYLATIONOFDNAMETHYLATIONPRO.
SUMOYLATIONOFSUMOYLATIONPROTEI.
MYD88 INDEPENDENT TLR4 CASCADE
DECTIN1MEDIATEDNONCANONICALNKS.
TCR SIGNALING
COMPLEX I BIOGENESIS
TICAM1RIP1MEDIATEDIKKCOMPLEXRE.
NEGATIVEREGULATIONOFNOTCH4SIGN.
DOWNREGULATIONOFERBB2ERBB3SIGN.
DISEASESOFCARBOHYDRATEMETABOLI.
ROSANDRNSPRODUCTIONINPHAGOCYTE.
PINK1 PRKN MEDIATED MITOPHAGY
REGULATIONOFHMOX1EXPRESSIONANA.
SIGNALINGBYTHEBCELLRECEPTORBCR.
THEACTIVATIONOFARYLSULFATASES.
INTERCONVERSIONOFNUCLEOTIDEDIT.
FORMATIONOFATPBYCHEMIOSMOTICCO.
GLYCOGEN STORAGE DISEASES
DEGRADATIONOFGLI1BYTHEPROTEASO.
IKKCOMPLEXRECRUITMENTMEDIATEBR.
NEDDYLATION
TP53 REGULATES METABOLIC GENES
INTERLEUKIN 1 SIGNALING
FCERIMEDIATEDNFKBACTIVATION.
METABOLISM OF POLYAMINES
CYTOPROTECTION BY HMOX1
ANTIGENPROCESSINGCROSSPRESENTA.
CRISTAE FORMATION
ASPARAGINENLINKEDGLYCOSYLATION.
GLYCOGEN SYNTHESIS
MITOCHONDRIAL PROTEIN IMPORT
PYRUVATEMETABOLISMANDCITRICATC.
PYRUVATE METABOLISM
PROTEIN LOCALIZATION
NEUTROPHIL DEGRANULATION
CELLULARRESPONSETOCHEMICALSTRE.
RESPIRATORY ELECTRON TRANSPORT
RRNA PROCESSING
RESPIRATORYELECTROTASBCCAHPBUP.
ACTIVATIONOFTHEMUBOTCBCAEASBT4.
SIGNALING BY ROBO RECEPTORS
INFLUENZA INFECTION
THECITRICACIDTCACYCLEANDRESPET.
NONSENSE MEDIATED DECAY NMD
REGULATIONOFEXPRESSIONOFSLITAR.
METABOLISMOFAMINOACIDSANDDERIV.
SELENOAMINO ACID METABOLISM
TRANSLATION
CELLULARRESPONSETOSTARVATION.
RESPONSEOFEIF2AK4GCN2TOAMINOAD.
EUKARYOTICTRANSLATIONELONGATIO.
EUKARYOTICTRANSLATIONINITIATIO.
SRPDEPENDENTCOTRANSLATIONAPTTM.
signed log10−scaled p.adjust

44

Figure S4

**Figure S4: Global differential expression and Reactome gene-set enrichment analysis.** **(A)** Left: Venn diagram of differentially expressed genes (DEGs) in female (fXO), male1 and male2 (mXO1, mXO2) 45,X vSMCs. Significance of pairwise overlapping DEGs shown by boldface p-values (hypergeometric distribution). Right: Differential vst heatmaps for overlapping DEG sets from the Venn diagram (left) as denoted by arrows. Ratio and p value (sign-test) denote the number of DEGs with concordant direction ("Dir.") in the triple and all pairwise overlapping DEG sets, with annotation panels showing the Wald statistic (DESeq2) for each gene. Dendrogram segregates samples by karyotype, irrespective of donor. **(B)** Reactome gene set enrichment analysis (GSEA), ordered by the quantile-normalized mean Wald statistic across 45,X conditions (ave.XO). The x axis denotes log-scaled GSEA-adjusted p value. Colors indicate normalized enrichment score of upregulated (red) and downregulated (blue) gene sets (sized by number of genes). Results for all isogenic monosomy-X comparisons (fXO, mXO1/2), averaged (ave.XO/ave.mXO), and control comparisons (m1m2E: male1/2 euploids; m1m2O: male1/2 45,X).

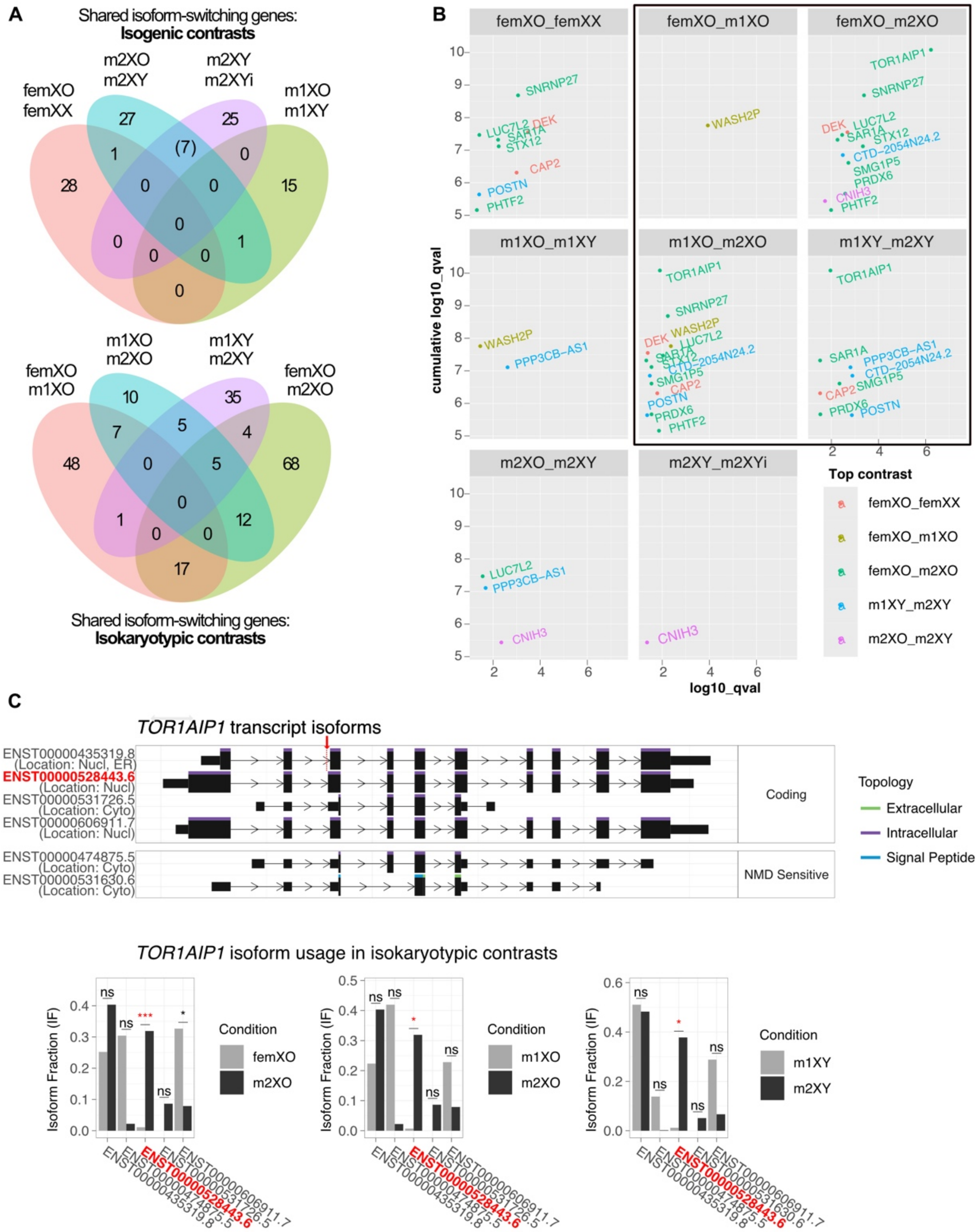
A
Shared isoform-switching genes:
Isogenic contrasts
femXO femXX
m2XO m2XY
m2XY m2XYi
m1XO m1XY
Shared isoform-switching genes:
Isokaryotypic contrasts
femXO m1XO
m1XO m2XO
m1XY m2XY
femXO m2XO
B
femXO_femXX
femXO_m1XO
femXO_m2XO
m1XO_m1XY
m1XO_m2XO
m1XY_m2XY
m2XO_m2XY
m2XY_m2XYi
cumulative log10_qval
log10_qval
Top contrast
femXO_femXX
femXO_m1XO
femXO_m2XO
m1XY_m2XY
m2XO_m2XY
C
TOR1AIP1 transcript isoforms
ENST00000435319.8 (Location: Nucl, ER)
ENST00000528443.6 (Location: Nucl)
ENST00000531726.5 (Location: Cyto)
ENST00000606911.7 (Location: Nucl)
ENST00000474875.5 (Location: Cyto)
ENST00000531630.6 (Location: Cyto)
Coding
NMD Sensitive
Topology
Extracellular
Intracellular
Signal Peptide
TOR1AIP1 isoform usage in isokaryotypic contrasts
Isoform Fraction (IF)
Condition
femXO
m2XO
m1XO
m1XY
m2XY

**Figure S5**

**Figure S5: Transcript isoform usage in isogenic and isokaryotypic contrasts.**

**(A)** Overlap in genes with transcript significant isoform switches (q-value ≤ 0.05, minimal difference of 20%) shared among (*top)*: isogenic contrasts (within each donor), or (*bottom)*: isokaryotypic contrasts (across donors). Contrasted conditions indicated above each Venn diagram. **(B)** Genes with transcript isoform switches detected in 3 or more contrasts plotted with cumulative log10-scaled q-value (sum of FDR-adjusted p values across all significant contrasts) over contrast-specific q-value in isogenic and isokaryotypic contrasts (boxed), with the highest scoring q-value contrast denoted by color (top_contrast). **(C)** Alternative splicing site acceptor lengthening exon 3 of *TOR1AIP1* (*top,* coordinates not to scale*),* previously reported as a lead splicing QTL in cardiovascular disease, is significantly increased in all isokaryotypic contrasts involving male2 (*bottom*).

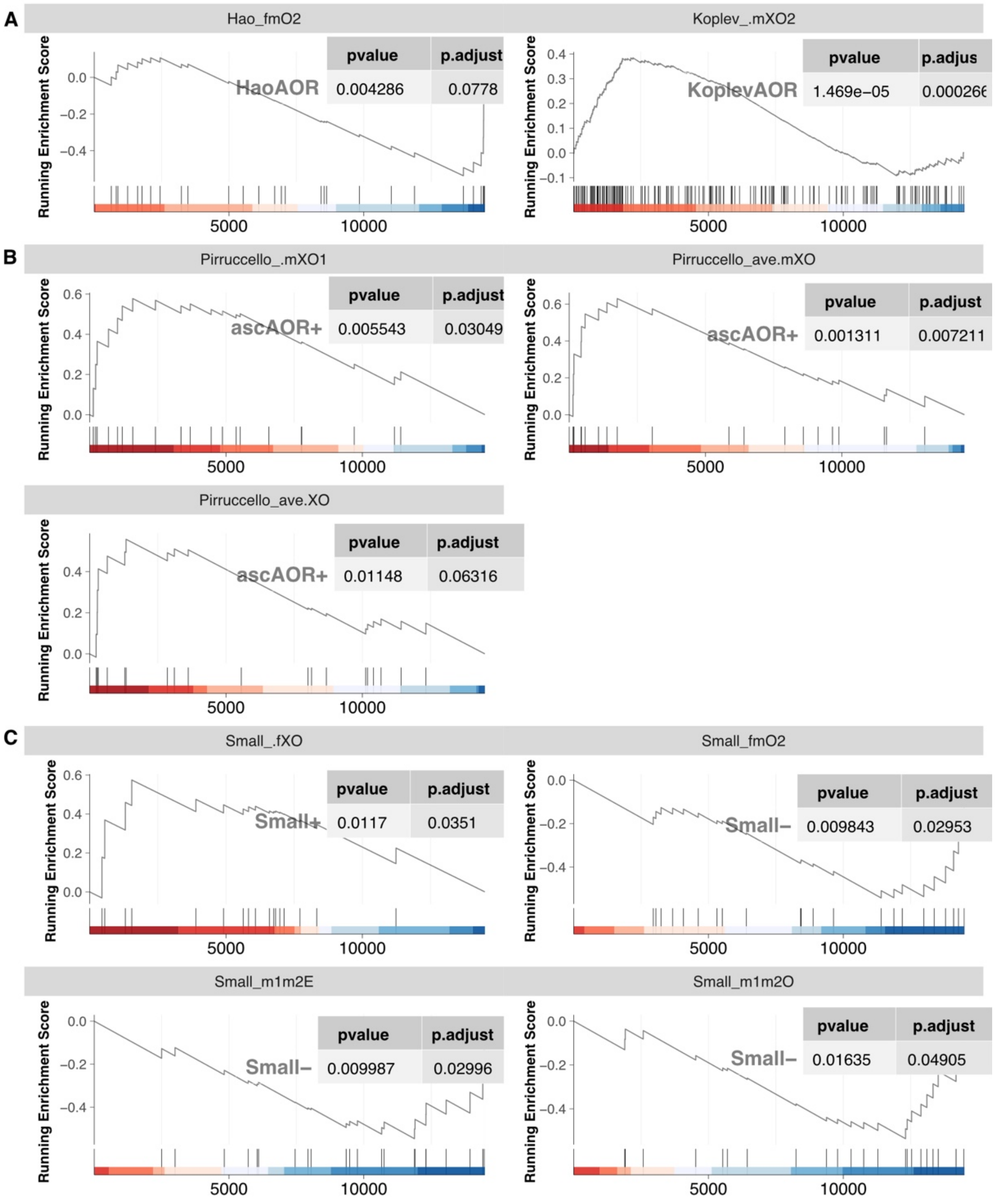


**Figure S6: GSEA plots of gene sets associated with cardiovascular disease or traits.**

Plots are faceted by study and the enriched condition. GSEA running enrichment score (y-axis) is plotted over genes (x-axis) by descending Wald statistic (DESeq2). Enriched study-category gene set (HaoAOR, KoplevAOR, ascAOR+, and Small-/+ denoted in inset table listing nominal and adjusted fgsea p-values.
